# The actin binding protein α-adducin modulates desmosomal turnover and plasticity

**DOI:** 10.1101/787846

**Authors:** Matthias Hiermaier, Felix Kliewe, Camilla Schinner, Chiara Stüdle, I. Piotr Maly, Marie-Therès Wanuske, Vera Rötzer, Nicole Endlich, Franziska Vielmuth, Jens Waschke, Volker Spindler

**Author notes:** **Address for Correspondence:** Dr. Volker Spindler Department of Biomedicine, Institute of Anatomy University of Basel Pestalozzistrasse 20 4056 Basel, Switzerland.

## Abstract

Intercellular adhesion is essential for tissue integrity and homeostasis. Desmosomes are especially abundant in the epidermis and the myocardium, tissues, which are under constantly changing mechanical stresses. Yet, it is largely unclear whether desmosomal adhesion can be rapidly adapted to changing demands and the mechanisms underlying desmosome turnover are only partially understood. We here show that loss of the actin-binding protein α-adducin prevented the ability of cultured keratinocytes or murine epidermis to withstand mechanical stress paralleled with reduced desmosome number. This effect was not caused by decreased levels or impaired adhesive properties of desmosomal molecules but rather by altered desmosome turnover. Mechanistically, reduced cortical actin density in α-adducin knockout keratinocytes resulted in increased mobility of the desmosomal adhesion molecule desmoglein 3 (Dsg3) and impaired interactions with E-cadherin, a crucial step in desmosome formation. Accordingly, loss of α-adducin prevented increased membrane localization of Dsg3 in response to cyclic stretch or shear stress. Our data demonstrate plasticity of desmosomal molecules in response to mechanical stimuli and unravel a mechanism how the actin cytoskeleton indirectly shapes intercellular adhesion by restricting the mobility of desmosomal molecules.

## INTRODUCTION

All multicellular organisms rely on cell-cell and cell-matrix interactions to facilitate the integrity of tissues and to sense and adapt to changes within the environment. These interactions are dependent on membrane receptors and adhesion molecules. The latter are often found in ultrastructurally detectable clusters linked to the cytoskeleton, which are then termed cell junctions. Desmosomes are an important type of junctions linking two adjacent cells, being present in all epithelial and some non-epithelial tissues. The desmosomal cadherins desmogleins (Dsg) 1-4 and desmocollins 1-3 are transmembrane molecules and build the core of desmosomes. The extracellular domains mediate adhesion in a calcium-dependent manner (Delva et al., 2009, Saito et al., 2012, Thomason et al., 2010) whereas the intracellular domains couple to proteins of the armadillo family (plakoglobin and plakophilin 1-3) (Carnahan et al., 2010, Hatzfeld, 2007). These in turn are linked to desmoplakin (Bornslaeger et al., 1996, Gallicano et al., 1998, Sonnenberg and Liem, 2007), which connects the whole protein complex to the intermediate filament cytoskeleton. Due to their biochemical stability, desmosomes were long considered to be largely static complexes, but have meanwhile been proven to be dynamic structures undergoing constant remodeling even under steady state conditions (Nekrasova and Green, 2013, Windoffer et al., 2002). Specifically, adhesion molecules can locate diffusibly in the membrane outside of desmosomes, which may represent a pool for the shuttle of desmosomal cadherins into or out of the desmosome. Moreover, desmosomes can maturate over time, leading to junctional complexes, which are largely resistant to disassembly in response to reduced extracellular Ca^2+^ levels (Garrod and Kimura, 2008). Desmosomes were reported to locate to cholesterol-rich membrane domains (lipid rafts) (Lewis et al., 2019, Stahley et al., 2014) and desmosome assembly is dependent on actin cytoskeleton remodeling (Godsel et al., 2010). The detailed mechanisms how desmosomal cadherins are targeted to desmosomes, however, are only partially understood.

We have recently shown that loss of the actin-binding protein α-adducin leads to reduced intercellular adhesion (Roetzer et al., 2014). Of the three adducins, α- and γ-adducin are ubiquitously expressed (Dong et al., 1995, Gilligan et al., 1999, Gilligan et al., 2002, Joshi et al., 1991). They form heterodimers and heterotetramers (Hughes and Bennett, 1995) which is important for their functionality (Gardner and Bennett, 1986, Li et al., 1998). Adducins are crucial in organizing the spectrin-actin meshwork underneath the cell membrane, specifically by bundling of actin filaments (Mische et al., 1987, Taylor and Taylor, 1994), capping of the fast growing ends (Kuhlman et al., 1996) and by recruiting spectrin to actin filaments (Bennett et al., 1988, Gardner and Bennett, 1987, Hughes and Bennett, 1995). Adducins are phosphorylated in response to RhoA signaling, which can be exploited to mitigate adhesion loss in pemphigus, an autoimmune blistering skin disease caused by autoantibodies against Dsg3 and Dsg1 (Roetzer et al., 2014, Waschke et al., 2006). Given the link between desmosomal adhesion and α-adducin, we here studied the underlying mechanisms and the impact of α-adducin on desmosome plasticity and turnover.

## RESULTS

### Loss of α-adducin Leads to Decreased Intercellular Adhesion

To understand the role of α-adducin for desmosome function, we established two α-adducin knockout (ko) cell culture models. First, CRISPR/Cas9 knockouts were generated in the human HaCaT keratinocyte cell line and compared to corresponding control cell lines (ctrl). Second, we isolated and immortalized primary murine keratinocytes (MEK) from α-adducin ko mice and corresponding wildtype (wt) littermates. The deletion of α-adducin was confirmed by Western blotting and immunostaining (Sup. Fig. 1A-C). Quantification of Western blot bands showed no significantly altered expression of desmosomal proteins in both models that were consistent throughout all clones. Similarly, the adherens junction’s protein E-Cadherin (E-Cad) was unaltered. However, as tested in dispase-based dissociation assays, all knockout clones independent from the background showed severely compromised abilities to withstand mechanical stress. (Fig. 1A, B). Similar results were observed in a modification of the dissociation assay using epidermal biopsies from α-adducin (Add1) ko and wt fetuses harvested at E18.5 (Fig. 1C), demonstrating that knockout skin is less resilient to mechanical stress. The skin of these mice showed no altered epidermal thickness or changes in Dsg3 localization and abundance, respectively (Sup. Fig. 1D, E). For further experiments, we selected the HaCaT ctrl clone 2 and the ko clone 2 based on their morphological similarities with the parental HaCaT line. To confirm whether indeed desmosomes were affected by α-adducin loss, we performed transmission electron microscopy (Fig. 1D, E, Sup. Fig. 1F). While no alterations in desmosome length were evident, α-adducin ko HaCaT cells exhibited significantly less desmosomes compared to ctrl cells. Furthermore, the morphology of desmosomes was different, with a blurrier appearance of the extracellular space in ko cells and less defined intracellular plaque regions. These results outline the importance of α-adducin for intercellular adhesion and resistance to mechanical stress and suggest a dysregulation of desmosomes on a level independent of protein expression.

**Fig. 1.**
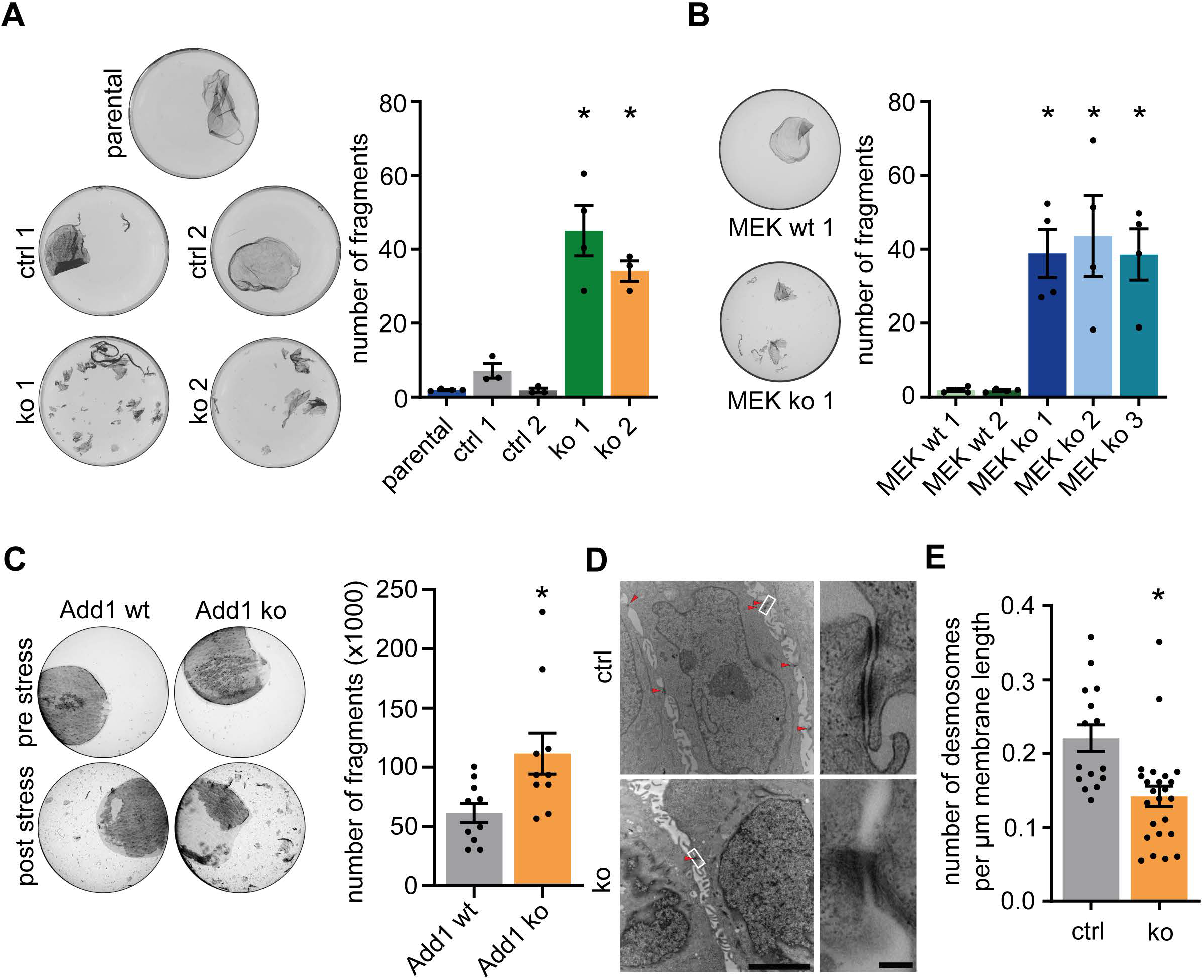
Loss of α-adducin impairs resilience to mechanical stress and reduces desmosome numbers. (**A**) Representative images of monolayer fragments after application of shear stress and quantification of dispase-based dissociation assays of parental HaCaT, α-adducin ctrl and ko cells (left). n≥3, each dot represents the mean value of 3 replicates of one independent experiment, * p<0.05 vs. parental, error bars represent SEM. (**B**) Representative images of monolayer fragments after application of shear stress and quantification of dispase-based dissociation assays of wt MEK cell lines and α-adducin ko MEK cell lines. n=4, each dot represents the mean value of 3 replicates of one independent experiment, * p<0.05 vs. MEK wt 1, error bars represent SEM. (**C**) Representative images of murine skin biopsies in dispase-based dissociation assays. Whereas the Stratum corneum (lightly stained) is unaffected, the lower cell layers (strongly stained) dissociate in response to shear stress. After application of rotating stress, cell fragments released from biopsy were quantified in a hemocytometer. n=10 mice per genotype, each dot represents one animal, * p<0.05 vs. Add1 wt, error bars represent SEM. (**D**) Representative electron microscopic images of confluent HaCaT α-adducin ctrl and ko cells. Desmosomes are marked by red arrowheads. White rectangles mark the zoomed area on the right. Scale bar overview: 5 µm. Scale bar zoom: 250 nm. (**E**) Quantification of desmosome density corresponding to **D**. n≥15 cells, each dot represents the value of one cell, * p<0.05 vs. ctrl, error bars represent SEM.

### Loss of α-adducin Reduces Cytoskeletal Anchorage of Desmosomal Cadherins

We have previously shown that the interaction forces of single desmosomal cadherins depend on signaling cues from within the cell and the connection to structural molecules such as keratins (Vielmuth et al., 2018b). To test whether the reduction of intercellular adhesion in α-adducin ko cells was due to a decreased interaction force between desmosomal cadherins, atomic force microscopy (AFM) single molecule force spectroscopy experiments were conducted. Cantilevers functionalized with recombinant Dsg3 extracellular domains were used to generate adhesion maps on the surface of living HaCaT cells as previously described (Vielmuth et al., 2018a). Incubation with a monoclonal antibody against the outermost extracellular domain reduced the binding probability and served as specificity control (Sup. Fig. 2A). The resulting force-distance curves containing interaction events were further classified depending on their appearance (Fig. 2A). Either they showed an increasing negative slope just before the rupture of the bond (“bent” unbinding event) or a flat slope, indicating a constant force acting on the cantilever despite increasing distance between cell surface and AFM head (“plateau”). As described earlier (Friedrichs et al., 2013), the former corresponds to a binding partner being rigidly anchored to the cytoskeleton whereas the latter indicates a non-anchored interacting molecule which is pulled away from the cell together with the surrounding membrane, resulting in the formation of a membrane tether. Deficiency of α-adducin did not lead to changes of the overall interaction probability (Fig. 2B) and interaction forces (Fig. 2C) between the Dsg3 extracellular domain on the cantilever and its binding partner on the cell membrane. However, when separating unbinding events in bent and plateau, we detected a shift of the interaction probability from the anchored to the non-anchored type of interaction in α-adducin ko cells (Fig. 2D). This shift towards tether unbinding events was also visible when plotting the unbinding forces and their corresponding step distances in a heat map depicting their relative probability in the force-distance space (Fig. 2E). These data indicate a reduced cytoskeletal anchorage of desmosomal cadherins in the membrane but did not support the hypothesis that reduced interaction strength on a single molecule level causes the adhesion-deficit of α-adducin ko cells.

**Fig. 2.**
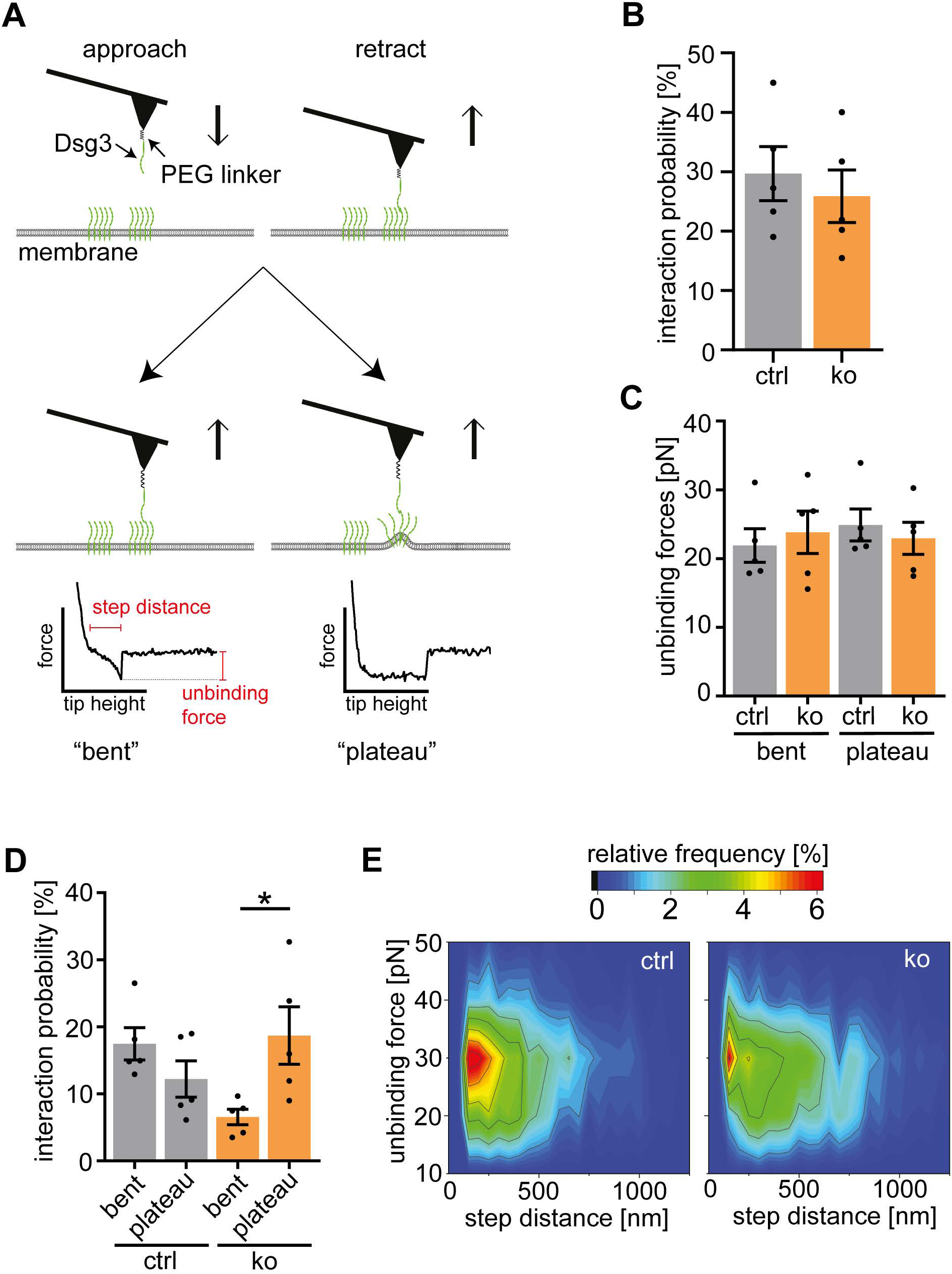
Loss of α-adducin does not change interaction forces of Dsg3 molecules. (**A**) Schematic of the two types of unbinding events detected by AFM. (**B-E**) AFM force mapping of Dsg3-functionalized cantilevers probed on living ctrl and α-adducin ko HaCaT cells. Analysis of (**B**) total interaction probability, (**C**) unbinding forces and (**D**) interaction probability, divided into bent and plateau unbinding events as shown in **A**, n=5, each dot represents the mean value of >2000 force-distance-curves from 5 independent experiments, error bars represent SEM. * p<0.05. (**E**) Heat map of the relative frequency of Dsg3 interaction events plotted as force-distance map. n>1500 interaction events from 5 independent experiments.

### Loss of α-adducin Accelerates Dsg3 Turnover During Desmosome Assembly and Disassembly

As other possibility how α-adducin influences desmosomes, we next tested whether altered desmosome dynamics correlated with the reduced adhesion observed in α-adducin ko. The Ca^2+^ dependence of desmosomes can be used to trigger desmosome disassembly by removing extracellular Ca^2+^ or to prime desmosome assembly by rising extracellular Ca^2+^ levels (Jones and Goldman, 1985). During desmosome disassembly, immunostainings of Dsg3 revealed reduced levels of Dsg3 in the cell membrane of α-adducin ko compared to baseline conditions (Fig. 3A, B). In addition, the remaining Dsg3 signal appeared more fragmented along membranes (Sup. Fig. 2B), indicating a more inhomogeneous depletion of Dsg3 from the cell membrane. Cell surface molecule biotinylation experiments supported these results, showing significantly reduced membrane-bound Dsg3 in α-adducin-deficient cells after 1 h Ca^2+^ depletion, while E-Cad was not altered (Fig. 3C). Induction of desmosome assembly through 4 h Ca^2+^ repletion led to faster recruitment of Dsg3 to the cell membrane in α-adducin-deficient cells (Fig. 3A, D). However, this faster recruitment of Dsg3 did not correlate with increased adhesion, as α-adducin ko cells showed a slower decrease of fragment numbers over Ca^2+^ repletion time (Fig. 3E). 24 h after Ca^2+^ repletion, α-adducin ctrl cells had reached their regular level of adhesion whereas α-adducin ko cells still showed significantly more fragments compared to non-switched cells. Recent data indicate that desmosome maturation, finally resulting in Ca^2+^ independency, correlates with reduced turnover of desmosomal cadherins (Bartle et al., 2019). The results from the immunostaining experiments und Ca^2+^ depletion conditions (Fig. 3A, B) already suggest reduced resistance to Ca^2+^ removal. Indeed, chelation of Ca^2+^ for 90 min as suggested to test Ca^2+^ independency (Garrod et al., 2005) led to increased fragmentation in α-adducin ko cells, demonstrating reduced desmosome maturation in response to α-adducin loss (Fig 3F).

**Fig. 3.**
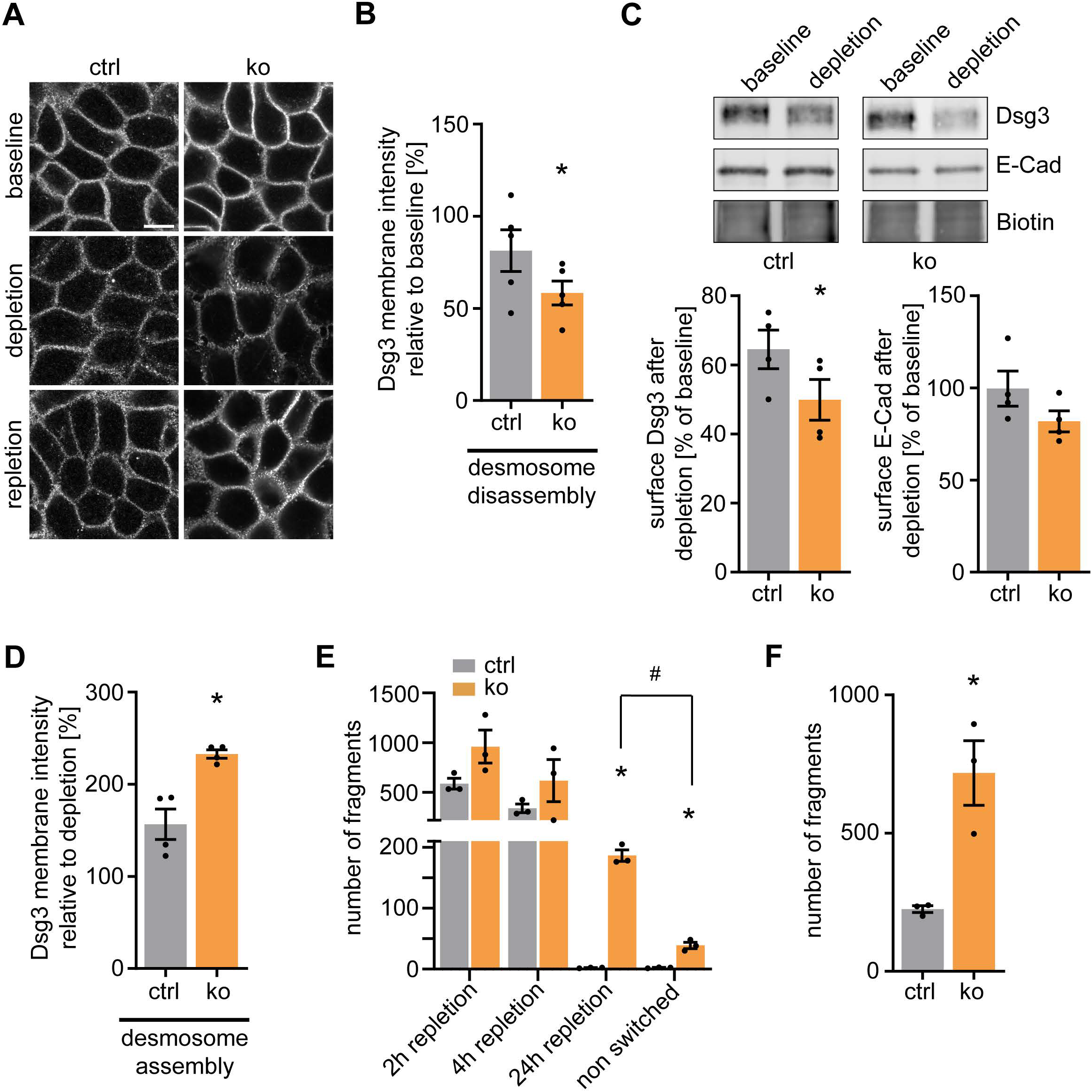
Loss of α-adducin accelerates Dsg3 turnover during desmosome assembly and disassembly. (**A**) Representative immunostainings of α-adducin ctrl and ko HaCaT cells before (baseline), after 1 h of Ca^2+^ depletion (depletion) and after 4 h of Ca^2+^ repletion following depletion (repletion), stained for Dsg3. Scale bar: 10 µm. (**B**) Quantification of Dsg3 fluorescence intensity along the membrane during desmosome disassembly (relative to baseline). n=5, each dot represents the mean of >6 cells of one independent experiment, * p<0.05 vs. ctrl, error bars represent SEM. (**C**) Western blot analysis after isolation of cell surface molecules via biotinylation in α-adducin ctrl and ko HaCaT cell lysates before (baseline) and after 1 h Ca^2+^ depletion probed for Dsg3 and E-Cad with biotin as a loading control (top). Respective densitometric quantification of Western blots (bottom). n=4, each dot represents one densitometric measurement of one independent experiment, * p<0.05 vs. ctrl, error bars represent SEM. (**D**) Quantification of immunostainings of Dsg3 fluorescence intensity along the membrane after 4 h of Ca^2+^-induced desmosome assembly (relative to depletion). n=4, each dot represents the mean of >6 cells of one independent experiment, * p<0.05 vs. ctrl, error bars represent SEM. (**E**) Quantification of fragment numbers in dispase-based dissociation assays of α-adducin ctrl and ko HaCaT cells without Ca^2+^ switch and after 2 h, 4 h and 24 h of Ca^2+^ repletion, respectively. n=3, each dot represents the mean of 2 replicates of one independent experiment, * p<0.05 ko vs. respective ctrl; # p<0.05, error bars represent SEM. (**F**) Quantification of fragment numbers in Ca^2+^-independency assays of α-adducin ctrl and ko HaCaT cells. n=3, each dot represents the mean of ≥3 replicates of one independent experiment, * p<0.05 vs. ctrl, error bars represent SEM.

Taken together these results suggest increased Dsg3 membrane turnover in α-adducin ko cells. However, this was not paralleled with increased intercellular adhesion, indicating dysfunctional desmosome assembly.

### Loss of α-adducin Enhances Dsg3 Mobility in the Cell Membrane

To analyze the increased turnover of Dsg3 in more detail, we conducted fluorescence recovery after photobleaching (FRAP) experiments. After bleaching of Dsg3-GFP signals at the cell-cell contact area of two adjacent cells, α-adducin ko cells showed a faster recovery of fluorescence intensity than control cells and a larger pool of mobile Dsg3-GFP (Fig. 4A, B). This observation may be explained either by increased transport of Dsg3-GFP from inside the cell to the cell membrane or by increased lateral mobility within the cell membrane. To test the first hypothesis, Dsg3-GFP signals were tracked from a perinuclear region towards the cell border. Analysis of the resulting trajectories showed no significant changes in movement speed and only a minor reduction in trajectory linearity (Sup. Fig. 3A, B). Thus, we investigated movement of Dsg3 molecules in the cell membrane. To visualize Dsg3 on the cell surface, we functionalized quantum dots with single chain variable fragments (scFv) directed against Dsg3 derived from a phage display approach of a pemphigus patient (Payne et al., 2005). The use of a non-pathogenic anti-Dsg3-scFv fraction ensured that Dsg3 was not internalized upon binding in contrast to a pathogenic anti-Dsg3 scFv (Sup. Fig. 3C) or complete anti-Dsg3 antibodies as reported in other studies (Schloegl et al., 2018). The movement of Dsg3 in control cells was mainly confined to an area of about 1 µm^2^, whereas in α-adducin-deficient cells it was much more spread out (Fig. 4C). In addition, Dsg3 speed in the cell membrane was significantly increased (Fig. 4D) and analysis of trajectory geometry demonstrated a strongly increased mean square displacement (Fig. 4E) in α-adducin ko cells. These results demonstrate that deficiency of α-adducin leads to increased movement of Dsg3 in the membrane, what possibly accounts for the higher FRAP in α-adducin-deficient cells.

**Fig. 4.**
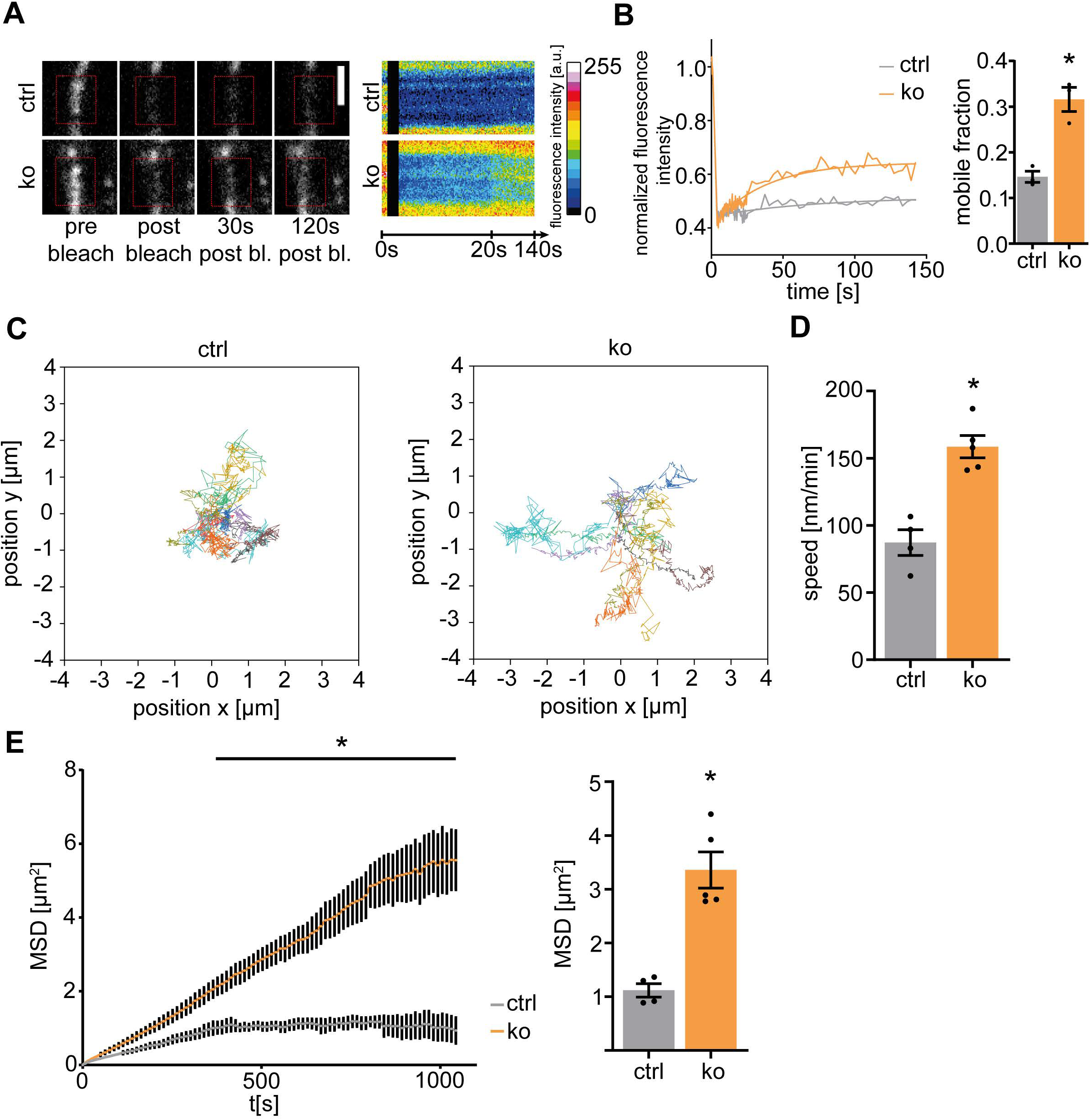
Loss of α-adducin enhances Dsg3 movement in the cell membrane. (**A)** Representative time series of FRAP experiments of Dsg3-GFP in membrane areas of α-adducin ctrl and ko HaCaT cells (left) with kymographs of bleached areas (right). Red rectangles mark bleached areas. The black area in the kymographs represents the bleaching interval. Scale bar: 2 µm. (**B**) Representative normalized fluorescence intensity plots with corresponding exponential fit for α-adducin ctrl and ko HaCaT cells in Dsg3-GFP FRAP experiments (left) and quantification of the average fraction of mobile molecules as calculated from fitted curves. n=3, each dot represents the mean of 4 FRAP experiments in 2 different cell pairs of one independent experiment, * p<0.05 vs. ctrl, error bars represent SEM. (**C**) Representative trajectories of anti-Dsg3-scFv coupled quantum dots on the surface of α-adducin ctrl and ko HaCaT cells. Temporal resolution is 4.2 s, total trajectory duration is 14 min. Corresponding quantification of (**D**) average quantum dot speed and (**E**) temporal development of the mean square displacement of quantum dot trajectories (left) and average mean square displacement after 10 min (right). n≥4, each dot represents the mean of >200 trajectories of one independent experiment, * p<0.05 vs. ctrl, error bars represent SEM.

### Loss of α-adducin Reduces the Interaction of Dsg3 with E-Cad During Desmosome Assembly

To elucidate potential mechanisms underlying the increased Dsg3 mobility in the cell membrane of α-adducin-deficient cells, we investigated the actin network underlying the membrane as cortical actin was shown to restrict the movement of membrane molecules (Sako et al., 1998, Wu et al., 2015). Structured illumination microscopy was used to obtain super-resolution images of the apical actin meshwork, which were subjected to skeletonization of filaments and further analysis (Fig. 5A). α-adducin-deficient cells exhibited a less dense and less connected apical actin network, evident by reduced number of filament junctions as well as reduced area underneath the cell membrane covered by actin filaments (Fig. 5B). Co-staining for Dsg3 revealed no differences in Dsg3 cluster size or intensity (Sup. Fig. 4A, B).

**Fig. 5.**
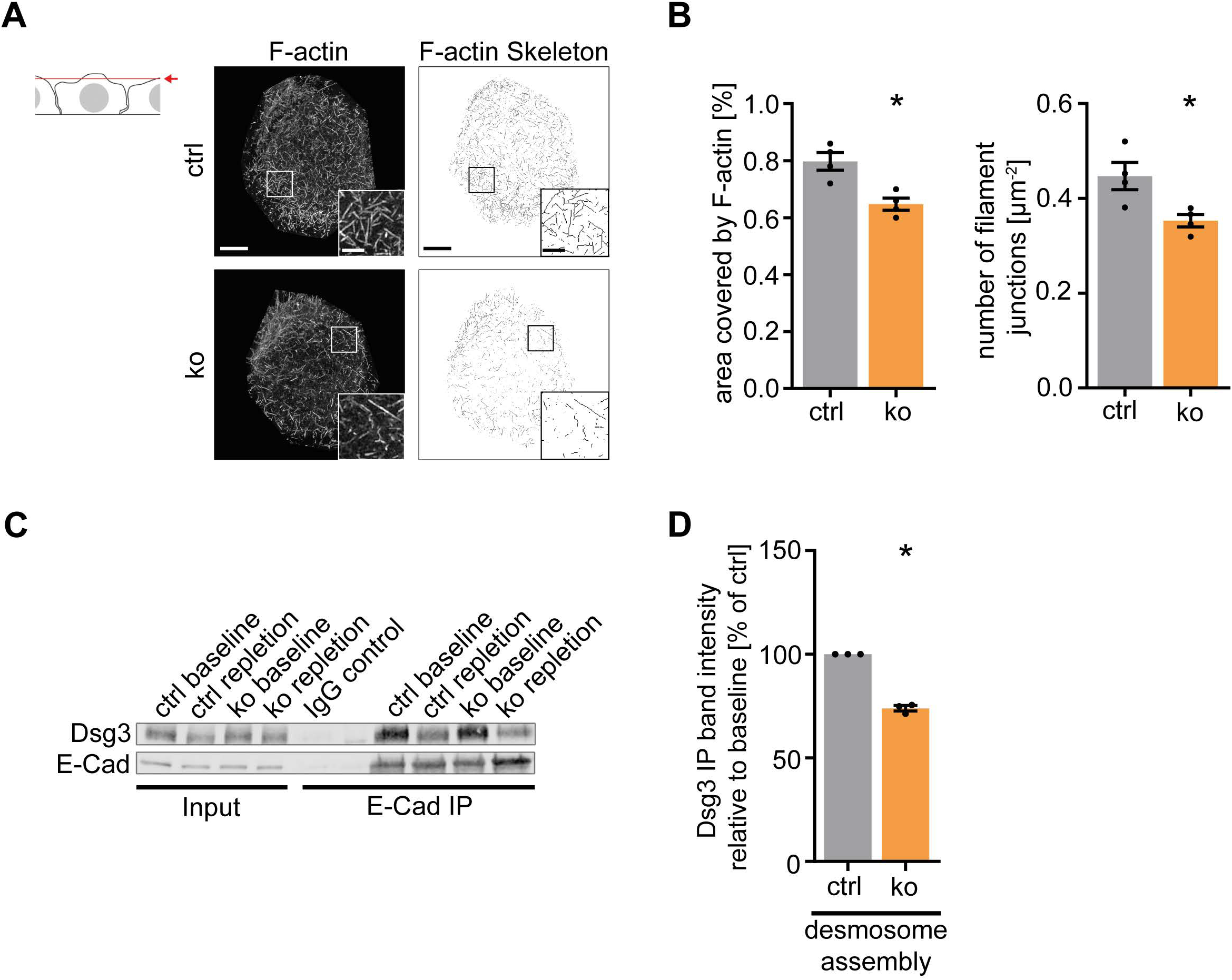
Loss of α-adducin decreases apical cortical actin meshwork density and reduces Dsg3/E-Cad interactions. (**A**) Schematic of the analyzed apical cell portion. Z-stacks comprising the apical 1.5 µm of a cell were maximum-intensity-projected in one plain and then further analyzed (upper left). Apical parts of α-adducin ctrl and ko HaCaT cells stained for F-actin (center) and corresponding skeletonization of actin filaments (right). Scale bar: 5 µm. Scale bar insert: 2 µm. (**B**) Quantification of F-actin skeletons of α-adducin ctrl and ko HaCaT cells. Percentage of the F-actin-covered area of the apical cell portion (left) and quantification of filament junctions per area (right). n=4, each dot represents the mean of >8 cells of one independent experiment, * p<0.05 vs. ctrl, error bars represent SEM. (**C**) E-Cad immunoprecipitations of α-adducin ctrl and ko HaCaT cells without (baseline) and after 1 h Ca^2+^ depletion and 4 h repletion (repletion). (**D**) Densitometry of Dsg3 bands in E-Cad immunoprecipitations after Ca^2+^ switch. n=3, each dot represents one densitometric measurement of one independent experiment, * p<0.05 vs. ctrl, error bars represent SEM.

It was shown that interactions of Dsg3 with E-Cad are required to translocate Dsgs in the cell membrane to form desmosomal precursors (Roetzer et al., 2015, Shafraz et al., 2018). To study whether this mechanism is dependent on α-adducin, we immunoprecipitated E-Cad during Ca^2+^-induced desmosome assembly. In α-adducin ko cells, a significant reduction of E-Cad/Dsg3 interactions was observed after 2 h of Ca^2+^ repletion compared to ctrl (Fig. 5C, D). Together, these data show that loss of α-adducin leads to a disturbed and less branched actin meshwork underneath the cell membrane and impairs the interaction of Dsg3 with E-Cad. These observations are consistent with a model in which under physiological conditions cortical actin restricts Dsg3 movement and thereby fosters Dsg3/E-Cad interaction, which is essential for desmosome formation.

### Loss of α-adducin Impairs Dsg3 Plasticity

Desmosomes, like other cell junctions, are constantly remodeled (Green et al., 2010). We hypothesized that this remodeling is necessary to adapt cell-cell adhesion to changing cues exerted by the surrounding cells and may be dependent on α-adducin. To test this, confluent cells were exposed to cyclic stretch for 24 h. Immunostaining for Dsg3 revealed that α-adducin ctrl cells responded with increased Dsg3 content in the cell membrane (Fig. 6A). In contrast, cells lacking α-adducin failed to increase Dsg3 levels showing even reduced amounts of membrane Dsg3 (Fig. 6A, B). To further support these findings, we performed surface molecules biotinylation experiments after stretching (Fig. 6C, D). This assay revealed an increase of Dsg3, but not E-Cad, membrane content in ctrl cells, which was absent in ko cells. In a second approach, we subjected cells to a constant liquid flow for 24 h, which applied a shear stress of 6.69 dyn/cm^2^ on the cell surface. Similar to the application of cyclic stretch, Dsg3 intensity was increased in control cells after application of flow but was reduced in ko cells (Fig. 6E, F). These results demonstrate that keratinocytes lacking α-adducin are not able to adapt to external forces by increasing Dsg3 levels in the membrane.

**Fig. 6.**
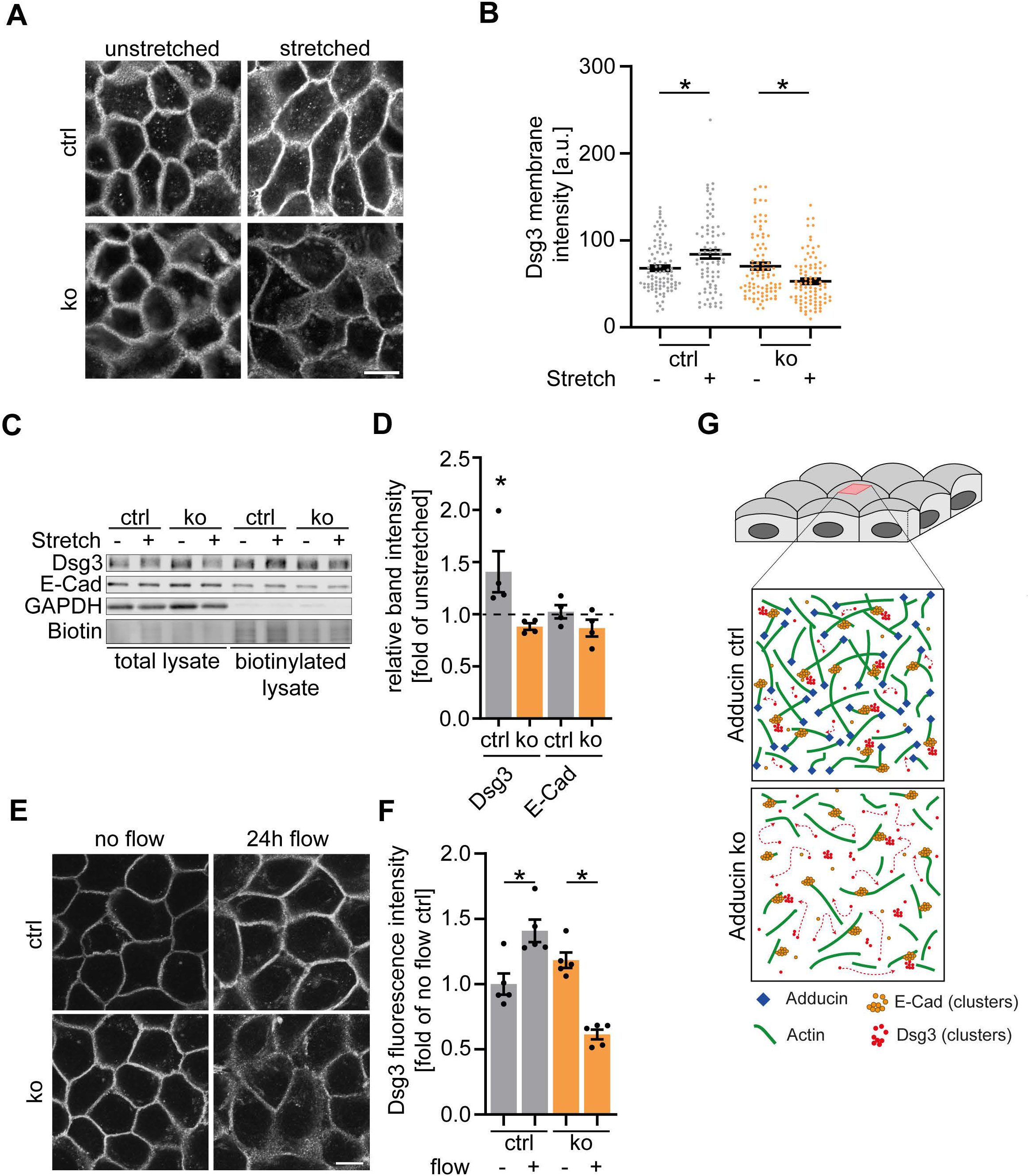
Dsg3 membrane levels are increased in response to external stress in α-adducin-dependent manner. (**A**) Representative immunostainings of Dsg3 of α-adducin ctrl and ko HaCaT cells subjected to 24 h of cyclic stretch. Scale bar: 20 µm. (**B**) Corresponding quantification of Dsg3 membrane intensity of immunostainings with or without stretch across the membrane between two cells. n= 90 cells of 3 independent experiments, * p<0.05, error bars represent SEM (**C**) Representative Western blot of surface molecule biotinylation assay of α-adducin ctrl and ko HaCaT cells subjected to either 24 h stretching or no stretching. (**D**) Corresponding densitometric analysis of Dsg3 and E-Cad bands of stretched relative to unstretched conditions. All bands were normalized to biotin bands. n=4, each dot represents one densitometric measurement of one independent experiment, * p<0.05 vs. unstretched, error bars represent SEM. (**E**) Representative immunostainings of α-adducin ctrl and ko HaCaT cell lines stained for Dsg3 with and without exposure for 24 h to laminar flow. Scale bar: 20 µm. (**F**) Corresponding quantifications of Dsg3 intensity along cell membranes. n=5, each dot represents the mean of >10 cells of one independent experiment, * p<0.05, error bars represent SEM. (**G**) Model of the effect of α-adducin loss on actin, E-Cad and Dsg3 during desmosome assembly.

## DISCUSSION

### Impact of α-adducin on Desmosome Formation

In this study, we demonstrate that α-adducin is required for proper desmosome assembly. The building bricks of desmosomes are assembled in distinct steps (Nekrasova and Green, 2013). Desmosomal cadherins are shuttled from the endoplasmatic reticulum to the membrane by microtubule-based transport. In agreement with this observation, we did not see profound changes during transport to the membrane if α-adducin, an actin-binding protein, was deleted. Recent work suggests that desmosomal cadherins undergo transient cis-interactions with E-Cad in the membrane during desmosome assembly (Roetzer et al., 2015, Shafraz et al., 2018). We show that loss of α-adducin results in a less dense cortical actin meshwork with increased mobility of Dsg3 molecules in the membrane which is paralleled with diminished interaction of Dsg3 with E-Cad. This indicates that the cortical actin restricts the movement of Dsg3 within the membrane in an α-adducin-dependent manner. Such a corralling function of the cortical actin was proposed more than two decades ago (Kusumi et al., 1999) and was recently revitalized for E-Cad by super resolution microscopy (Wu et al., 2015). Thus, we propose a model, in which the corralling of Dsg3 and presumably of E-Cad is necessary to efficiently meet each other for the transient interactions taking place during desmosome formation (Fig. 6G). The reduced actin meshwork density in α-adducin ko cells allows greater movement of Dsg3 and, potentially, E-Cad, which impairs these processes. Our observation by TEM of reduced desmosome numbers in α-adducin-deficient cells supports this hypothesis.

Nascent desmosomes serve as nucleation sites for keratin filaments (Godsel et al., 2005, Moch et al., 2019). Vice versa, the binding properties of desmosomal cadherins are compromised if keratins are absent, indicating an inside-out regulation of desmosomal adhesion on the single molecule level by intermediate filaments (Vielmuth et al., 2018b). However, since we did not find any alterations in the binding force of Dsg3 molecules on α-adducin ko cells, these data do not favor a model in which the cortical actin cytoskeleton shapes adhesion on a single molecule level.

Additionally, the desmosomal complex locates to lipid rafts, although the precise mechanisms here are only partially elucidated. Recent work outlined that the transmembrane domain of desmosomal cadherins is larger than that of classical cadherins, which leads to a preference of desmosomal cadherins to locate within the slightly thicker cell membrane present in lipid rafts (Lewis et al., 2019). Interestingly, β-adducin was shown to locate to lipid rafts, too (Xu et al., 2015). As the attenuation of the Dsg3/E-Cad interaction in α-adducin ko cells was only partial, it is possible that α-adducin is also required to sequester desmosomal cadherins within lipid rafts. In line with this hypothesis, the increase of interactions with non-cytoskeletal desmosomal cadherins as indicated by our AFM force spectroscopy experiments may point to a larger pool of “free” molecules outside of lipid rafts.

### α-Adducin Shapes Plasticity of Desmosomes

It is well established that desmosomal adhesion is not uniform but can switch between different maturation states. For instance, culturing cells for long time periods of confluency leads to Ca^2+^ independency (Kimura et al., 2007). Under wound healing conditions and dependent on PKC signaling, this mature state is reverted and desmosomes regain Ca^2+^ dependency (Garrod et al., 2005). In α-adducin ko cells, we also noted increased sensitivity to Ca^2+^ depletion, which may be a result of increased Dsg turnover (Bartle et al., 2019). We reasoned that in addition to these two extreme states, i.e. Ca^2+^ dependency and independency, desmosomes also need to be able to flexibly adapt adhesion to external stimuli. For instance, the epidermis is under constantly changing load due to friction or pressure. Under certain circumstances, e.g. exercise, it is conceivable that specific regions of the skin are subject to sustained above-average mechanical stresses, which also challenges intercellular junctions. These junctions, and in particular desmosomes with their high abundance in the epidermis, should be able to adapt to these additional stresses. Indeed, reports of a load-dependent adaption of intercellular adhesion come from cardiomyocytes. Upon increasing the contractility of the heart by sympathetic stimulation, presumably leading to increased load on cardiomyocyte junctions, a rapid recruitment of desmosomal molecules to the intercalated disc with a concurrent reinforcement of intercellular adhesion occurs (Schinner et al., 2017). Our results showing that Dsg3 is recruited to the cell membrane in response to cyclic stretch and laminar flow support these data and strengthen the concept of high desmosomal plasticity, which can be flexibly tuned to adapt to the actual demands. Importantly, these processes are impaired upon α-adducin loss, indicating that the altered Dsg turnover we observe is responsible for this phenomenon. Surprisingly, a high turnover appears not to be beneficial in this regard, as indicated by our observation that despite increased membrane localization of Dsg3 upon repletion of Ca^2+^, overall cell adhesion was still perturbed in α-adducin ko cells. This suggests that the transition of cell adhesion molecules from a free membrane pool into desmosomes or de novo clustering of nascent desmosomes is perturbed in these cells. Alternatively, because also desmosome disassembly by Ca^2+^ removal was increased in α-adducin ko cells, it is possible that the stability of Dsg3 within the desmosome is also impaired. Together, our data suggest that the cortical actin cytoskeleton contributes to desmosomal plasticity by coordinating the turnover of desmosomal molecules.

## ABBREVIATIONS

Add1: α-adducin
AFM: atomic force microscope
BSA: bovine serum albumin
ctrl: control
Dsg: desmoglein
ChIP: chromatin immunoprecipitation
E-Cad: cadherin
FDC: force-distance curve
FRAP: fluorescent recovery after photobleaching
ko: knockout
MEK: murine keratinocyte
scFv: single chain variable fragment
wt: wildtype

## MATERIALS AND METHODS

Please refer to the supplementary methods.

## ACKNOWLEDGEMENTS

We thank Dr. Aimee Payne, Department of Dermatology, University of Pennsylvania, Philadelphia, USA, for providing scFv constructs and Dr. Luane Peters, The Jackson Laboratory, Bar Harbor, ME, for providing the Add1 knockout strain. We are indebted to Elias Walter and Dr. Mariya Radeva, Institute of Anatomy, LMU Munich, for fruitful discussions. We are grateful to the Imaging Core facility (IMCF, University of Basel) and in particular Alexia Ferrand as well as the Center for Cellular Imaging and Nanoanalytics (C-CINA, University of Basel) for their excellent support. Excellent technical assistance by Anja Fuchs, Martina Hitzenbichler, Sabine Mühlsimer and Aude Zimmermann is gratefully acknowledged.

Funded by German Research Council SP1300/1-3 and SP1300/3-1 to VS and BMBF STOP-FSGS 01GM1901B to NE.

