## Supplementary files for "The actin binding protein α-adducin modulates desmosomal turnover and plasticity"

### MATERIALS AND METHODS

#### Cell Line Generation and Cell Culture

All cells were kept in a humidified atmosphere containing 5% $CO_2$ at 37°C in sterile cell culture flasks and well plates (Greiner, Kremsmünster, Austria). HaCaT α-adducin knock-out cell lines were generated using the CRISPR/Cas9 system. The pCMV-Cas9-GFP vector containing a CMV-driven CAS9 cassette linked to GFP through a self-cleaving 2A peptide as well as a U6 driven sgRNA targeting either exon 2 or 3 of the human *ADD1* gene (Target site: AATGGTGATTCTCGTGCGG and TAATGCCAATAGGCCTGTGGGG, respectively) was purchased from Sigma-Aldrich, St. Louis, MO, USA). The human keratinocyte cell line HaCaT was transiently transfected using Turbofect (Thermo Fisher Scientific, Waltham, MA, USA) according to the manufacturer's protocol. After 24 h, GFP-expressing cells were single cell-sorted in 96-well plates and surviving clones were checked for α-adducin expression via Western blot and finally sequenced. Two verified knockouts and two clones with non-successful knockout (serving as controls) were selected. HaCaT cells were cultured according to standard protocols in DMEM (Sigma Aldrich) containing 1.8 mM calcium, supplemented with 10% fetal bovine serum (Merck, Darmstadt, Germany), 50 µg/ml streptomycin sulfate (VWR, Radnor, PA, USA), 50 U/ml penicillin (VWR), 4 mM L-glutamine (Sigma Aldrich).

Murine keratinocytes (MEK) from ADD1 knockout mice and corresponding wildtype littermates were isolated from the epidermis and immortalized as followed: Neonatal mice were decapitated, washed 5 min each in PBS, 80% ethanol, 10% povidone-iodine and again PBS. The skin was peeled off under sterile conditions and placed over night at 4°C in dispase II solution (>2.4 U/ml dispase II in HBSS buffer, Sigma Aldrich) with 1:250 500x gentamicin/amphotericinB (CELLnTEC, Bern,

Switzerland). After 12 h, the epidermis was separated from the dermis and spread out in accutase (Sigma Aldrich) for 20 min at room temperature. Gentle agitation led to release of single cells from the epidermis which were collected and placed into cell culture dishes in 0.06 mM calcium murine keratinocyte medium (DMEM:Ham's F12 PAN-Biotech, Germany) supplemented with 10% fetal bovine serum (Merck), 50 µg/ml streptomycinsulfate (VWR), 50 U/ml penicillin (VWR), 58.8 pM epidermal growth factor (Invitrogen, Carlsbad, CA), 120 pM cholera toxin, 62.5 nM transferrin, 862 nM insulin, 1.1 µM hydrocortisone (all Sigma Aldrich), 62.5 µM gentamycin (CELLnTEC). Cells were kept in an incubator at 35°C with 5% $CO_2$, the medium was changed every three days and upon reaching confluency, cells were transferred into a new culture dish of the same size. After around 6 passages cells could be expanded to larger culture dishes and frozen for later use. 24 h before experiments were conducted, 1.8 mM calcium was added to the medium.

All cells were quarterly checked for mycoplasm infections using PCR and were proven negative and HaCaT cells were routinely authenticated by STR profiling.

**Animal Experiments**

All animal experiments were carried out according to the protocol approved by the Cantonal Veterinary office of Basel-Stadt (License number 2973). $\alpha$-adducin transgenic animals (B6.129P2-Add1<tm1>/J) were described before (Robledo et al., 2008) and a kind gift of Dr. Luane Peters, The Jackson laboratory, Bar Harbor, ME, USA. Animals were bred heterozygously and were housed under SPF conditions according to institutional guidelines. Offspring were genotyped as outlined before (Robledo et al., 2008). Ko animals died in most cases within 24 h after birth due to unknown reasons. For harvesting of skin samples, we thus had to rely on ex utero dissections at E18.5, with the exception of some newborn animals that could be used to isolate MEKs at P1.

### Western Blot Analysis

Western blot analysis was performed according to standard procedures (Hartlieb et al., 2013). Briefly, cells were cultured in 24-well plates until they reached confluence. Cells were washed once with room temperature PBS and then scraped in lysis buffer containing 12.5 mM HEPES (Sigma Aldrich), 1 mM EDTA (VWR), 12.5 mM sodium fluoride (VWR), 0.5% sodium dodecyl sulfate (Sigma Aldrich) and a protease inhibitor cocktail according to the manufacturer's protocol (cOmplete, #11697498001, Roche Diagnostics, Mannheim, Germany) (pH 7.4). Subsequently lysates were sonicated and a BCA protein assay kit (Thermo Fisher Scientific) was used to determine the protein concentration according to the manufacturer's protocol. Prior to loading on a SDS-PAGE gel for electrophoresis, lysates were mixed 2:1 with modified Laemmli buffer (Laemmli, 1970) and boiled 5 min at 95°C. After electrophoresis in electrophoresis buffer proteins were transferred on nitrocellulose membranes (Thermo Fisher Scientific) in blotting buffer. Membranes were dried and subsequently blocked in either 5% skim milk solution (Sigma Aldrich) in TRIS-buffered saline containing 0.05% tween 20 (Thermo Fisher Scientific), 5% BSA in TRIS-buffered saline containing 0.05% tween 20 (Thermo Fisher Scientific), or Odyssey blocking solution (#927-50000 Li-Cor, Lincoln, NE). Primary antibodies were incubated over night at 4°C: anti-GAPDH (for human cells, #sc-47724), anti-GAPDH (for murine cells, #sc-365062), anti- $\alpha$-adducin (#sc-33633), anti-desmoplakin (#sc-390975), anti-plakophilin 1 (#sc-33636 all Santa Cruz, Dallas, TX, USA), anti-plakoglobin (#61005 Progen, Heidelberg, Germany), anti-plakophilin 3 (#651113 Progen), anti-desmoglein 3 (EAP3816 Elabscience, Biozol, Eching, Germany), anti-desmoglein 2 (#61002 Progen), anti-E-cadherin (#610181 BD Biosciences, San Jose, CA, USA). Following secondary antibodies were applied: Goat anti-mouse-HRP (115-035-

068 Dianova, Hamburg, Germany), goat anti-rabbit-HRP (111-035-045 Dianova), goat anti-rabbit 800CW (925-32210 LiCor), goat anti-mouse 680RD (925-68071 LiCor). For protein detections either an ECL reaction system with self-made ECL solutions was used or direct imaging of fluorescent dyes was performed in an Odyssey FC imaging system (LiCor).

### Cell Surface Molecules Biotinylation Assay

Membrane-bound protein amounts were quantified using cell surface molecule biotinylation assays. Cells were grown to confluency and washed with ice-cold PBS. Afterwards, they were incubated for 60 min with 0.25 mM EZ-link sulfo-NHS-biotin (Thermo Fisher Scientific) in HBSS on ice. Cells were then subsequently washed three times with ice-cold 100 mM glycin in PBS and three times with ice-cold PBS. Cells where incubated for 15 min on ice with biotinylation lysis buffer [50 mM NaCl, 10 mM PIPES, 3 mM $MgCl_2$, 1% (v/v) Triton X-100, pH 6.8] and the lysate was centrifuged at 4°C for 5 min at 10,000x g. The protein concentration of the supernatant was determined using a BCA protein assay kit (Thermo Fisher Scientific) and equal amounts of protein were incubated with pre-cleared NeutrAvidin Agarose Resin (Thermo Fisher Scientific) overnight on a rocker. On the next day, beads were washed 3 times with biotinylation lysis buffer, then resuspended in Laemmli buffer and denatured at 98°C for 5 min. The whole amount was loaded on a 10% polyacrylamide gel and subjected to gel electrophoresis, blotting and visualization as described in *Western Blot Analysis*. Streptavidin-HRP (RPN1231 GE Healthcare, Chicago, IL, USA) served to detect biotinylated proteins.

### Immunoprecipitations

Immunoprecipitation was performed as previously described (Schinner et al., 2017). Briefly, cells were washed with ice-cold PBS and lysed in RIPA buffer [10 mM $Na_2HPO_4$, 150 mM NaCl, 1%(v/v) Triton X-100, 0.25% (v/v) sodium dodecyl sulfate, 1% (w/v) sodium deoxycholat, pH 7.2 freshly supplemented with protease inhibitors 1:1000 leupeptin, aprotinin, pepstatin and 1:100 phenylmethylsulphonyl fluoride] for 30 min on ice. Lysates were then homogenized by passing 10 times through a 20G and subsequently a 27G needle on ice and pelleted for 5 min at 4°C with 7700x *g*. Protein concentrations in the supernatant were measured using a BCA protein assay kit (Thermo Fisher Scientific) and equal amounts of protein were incubated with anti-desmoglein 3 (EAP3816 Elabscience), anti-E-cadherin (#610181 BD Biosciences), normal mouse IgG (#sc-2025 Santa Cruz) and normal rabbit IgG (#sc-2027, Santa Cruz) over night at 4°C. Afterwards, lysates were incubated with pre-cleared protein G Dynabeads (Thermo Fischer Scientific) for 1 h at 4°C. After several washing steps, the beads were re-suspended in Laemmli buffer and proteins denatured at 95°C for 10 min. The whole amount was then loaded on a 10% polyacrylamide gel and subjected to gel electrophoresis, blotting and visualization as described in *Western Blot Analysis.*

### Immunostaining

Cells were grown to confluency on glass cover slips, treated as indicated, washed twice with PBS, and fixed for 20 min with 4% paraformaldehyde (Thermo Fisher Scientific) in PBS. After washing 3 times with TBS, cells were permeabilized by incubating 0.5% Triton X-100 (Thermo Fisher Scientific) in TBS for 10 min. Subsequently, cells were washed 3 times with 0.1% Triton X-100 in TBS and blocked in 2% bovine serum albumin (BSA) (VWR) in 0.1% Triton X-100 in TBS for 60 min. Primary antibodies were diluted in 2% BSA in 0.1% Triton X-100 in TBS and incubated over night at 4°C. Afterwards, cells were washed 3 times with 0.1% Triton X-100 in TBS and appropriate secondary antibodies in 2% BSA in 0.1% Triton X-100 in TBS were incubated for 50 min at room temperature. DAPI was added for 10 min to visualize nuclei. Cells were subsequently washed 3 times in 0.1% Triton X-100 in PBS and 3 times in double distilled water and mounted with ProLong Diamond Antifade (Thermo Fisher). Samples were imaged using a SP5 inverted confocal microscope with a 63x HC PL APO NA=1.2 objective (both Leica, Mannheim, Germany), a LSM 710 inverted confocal microscope with a 63x PL APO NA=1.4 objective (Zeiss, Oberkochen, Germany) or for structured illumination microscopy on a DeltaVision OMX-Blaze (Version 4; Applied Precision, Issaquah, WA) using a 60x PL APO NA=1.42 oil immersion objective (Olympus, Tokyo, Japan). Images were analyzed using ImageJ software (NIH, Bethesda, MD, USA).

To assess the irregularity of membrane staining after calcium depletion, an irregularity score was devised. The intensity profile of the whole circumference of a cell was plotted using ImageJ (NIH) and the corresponding average fluorescence intensity $\langle I \rangle$ was calculated. Then for each pixel along the cell membrane it was determined if that pixel was brighter than the average $\langle I \rangle$, in which case this pixel gave a contribution of its intensity minus the average intensity. This was done for all pixels along the cell membrane and the single contributions were added up. The resulting value was then divided by the number of pixels with an above-average intensity to normalize the value for different cell sizes. Finally, the result was divided by the average fluorescence intensity $\langle I \rangle$ to normalize for differently strong staining. In that way the irregularity score solely is a measure for how dominant intensity differences are along the cell membrane, with an irregularity score of 0 meaning all pixels have the same intensity.

$$Irregularity\ Score = \frac{\sum_{i=I(i)>\langle I \rangle} I(i) - \langle I \rangle}{\langle I \rangle * \sum_{i=I(i)>\langle I \rangle} 1}$$

**Immunostainings and HE Stainings of Murine Biopsies**

Murine fetus biopsies at E18.5 were frozen in Tissue-Tek (Leica, Wetzlar, Germany) and 10 µm thick cryosections were made using a Cryo Star NX70 cryostat (Thermo Scientific). Air-dried cryosections were fixed in formaldehyde solution, 37 wt. % (Sigma Aldrich) stained with hematoxylin (Harris modified, J.T. Baker, Phillipsburg, NJ, USA) and 1 % Eosin, aqueous (RAL Diagnostics, Martillac, France) following standard protocols and mounted in Eukitt (Sigma-Aldrich). Images were acquired on a IX83 wide field microscope (Olympus) using a 10x PLN PH NA=0.25 objective (Zeiss). To determine the thickness of the epidermis 3-7 images of one HE stained section per fetus were analyzed in ImageJ (NIH) using the polygon selection and the fragment line tool. For immunofluorescent stainings air-dried cryosections were fixed with 3 % glyoxal and 20 % ethanol, pH 4, permeabilized with 0.2 % triton, 20 mM EDTA, pH 9.15, blocked with 2 % bovine serum albumin and stained overnight with rabbit anti-Dsg3 (EAP3816, Elabscience), followed by incubation with Cy3-conjugated anti-rabbit secondary antibody (Dianova), and sections were mounted in ProLong Diamond Antifade (Thermo Fisher).

**Fluorescence Recovery After Photobleaching (FRAP)**

FRAP studies were performed as described previously (Schinner et al., 2017). Briefly, cells were seeded in 8-well imaging chambers (Ibidi, Martinsried, Germany). The plasmid pEGFP-N1-Dsg3 encoding for murine Dsg3 was a kind gift from Yasushi Hanakawa (Ehime University School of Medicine, Japan) and transfected using Turbofect. 48 h after transfection, FRAP measurements were performed on a SP5 inverted confocal microscope with a 63x HC PL APO NA=1.2 objective (both Leica) at 37°C with 5% $CO_2$ and constant humidity using a cage incubator (OKOLAB, Burlingame, CA, USA). The measurements were carried out and analyzed with the FRAP wizard software (Leica). Regions of interest were defined along the cell membranes of two neighboring Dsg3-GFP transfected cells. After 5 frames of recording the pre-bleach intensity, the GFP signal was bleached using the 488-nm laser line at 100% transmission for 10 frames and recovery of the fluorescence was recorded for 80 frames, each frame with a duration of 0.266 s. Afterwards, 40 additional frames were captured every 3 s. The fraction of mobile molecules was determined by:

$$mobile\ fraction = \frac{I_{plateau} - I_0}{1 - I_0}$$

Here, $I_{plateau}$ is the plateau intensity which is reached after sufficient recovery time and $I_0$ is the minimal intensity that was achieved right after bleaching. The values of both intensities were normalized by dividing by the pre bleach intensity value.

**Dsg3-GFP Tracking**

To analyze intracellular Dsg3 transport, cells were transiently transfected to express Dsg3-GFP as described above. The same experimental setup as for FRAP measurements was used. Z-stacks with 1 µm spacing were acquired to cover the whole cell with a temporal resolution of about 9 frames per minute for 25 min. The resulting videos were processed using ICY software (de Chaumont et al., 2012) (Version 1.9) and in particular the spot detector and spot tracker plugins. The trajectories of intracellular vesicles containing Dsg3-GFP were analyzed with regard to speed and linearity of movement, the latter being the net distance traveled divided by the total distance covered.

**Intramembrane Tracking of Dsg3 Using Quantum Dots**

To recognize cell membrane bound Dsg3, CdSe/ZnS core-shell type quantum dots (Sigma Aldrich) were coupled to single chain variable fragments (scFv's) recognizing parts of the extracellular domain of Dsg3. The scFv's were a generous gift from Aimee S. Payne (University of Pennsylvania, Philadelphia, PA, USA). The coupling was achieved in a metal-affinity driven self-assembly as described earlier (Sapsford et al., 2007). Briefly, a solution of 500 nM CdSe/ZnS core-shell type quantum dots with 500 µM scFv in PBS was produced and gently agitated for 60 min at room temperature. This solution was applied 1:10 in imaging medium to confluent cells and after 2 h incubation time cells were gently washed twice with PBS to remove unbound quantum dots. After application of fresh imaging medium (DMEM$^{gfp}$-2, Evrogen, Moscow, Russia, supplemented with 10% fetal bovine serum (Merck), 50 µg/ml streptomycin sulfate (VWR), 50 U/ml penicillin (VWR)) cells were imaged on the setup described in section FRAP. Z-stacks were acquired to cover the top of the cells with a spacing of 0.5 µm and a temporal resolution of 4.2 s per frame for 250 frames. The resulting videos were analyzed for the trajectories of the quantum dots using ICY software as described in Dsg3-GFP tracking.

**Dispase-Based Dissociation Assays**

Cells were grown to confluency in 24-well plates and after washing with HBSS, cells were incubated with dispase II solution (>2.4 U/ml dispase II in HBSS, Sigma Aldrich) for 20 min at 37°C until the cell monolayer detached from the well bottom. Dispase II solution was carefully removed and replaced with fresh pre-warmed HBSS with 50 µg/ml 3-(4,5-dimethylthiazol-2-yl)-2,5-diphenyltetrazolium bromide for fragment visualization. The monolayers were then subjected to a defined mechanical stress by pipetting 10 times using an electric pipette (Eppendorf, Hamburg,

Germany). A stereo microscope (Olympus) and a SLR camera (Canon, Tokyo, Japan) were used to capture images of dissociated cell fragments. The number of fragments, which is an inverse measure of intercellular adhesion, was determined automatically using ImageJ (NIH). A variation of this assay was used to test for $Ca^{2+}$ independency: After detachment of the cell layer from the well bottom, dispase II solution was removed and cell layers were incubated for 90 minutes in cell culture medium containing 5 mM EGTA at 37°C in 5% $CO_2$ atmosphere. Cells were further processed as described above.

For dissociation assays of murine epidermis, α-adducin wt and ko murine fetuses were dissected at E18.5. Whole skin punch biopsies (8 mm diameter) were taken from the lower back of the animals and incubated in dispase II solution at 4°C overnight to separate epidermis from dermis. The epidermis was carefully transferred to 500 µl HBSS with 50 µg/ml 3-(4,5-dimethylthiazol-2-yl)-2,5-diphenyltetrazolium bromide and incubated at 37°C for 15 min. Epidermal biopsies were subsequently subjected to mechanical stress by defined orbital rotation at 300 rpm (SSM1, Stuart, St. Staffordshire, UK). Resulting fragments were collected, mixed and counted in a Neubauer hemocytometer. Total fragment number was calculated according to manufacturer's instructions.

**Atomic Force Microscopy (AFM)**

For AFM experiments, an atomic force microscope (Nanowizard IV, JPK Instruments, Berlin, Germany) mounted on an inverted microscope (IX83, Olympus) was used. AFM cantilevers (MLCT AFM Probes, Bruker, Calle Tecate, CA, USA) were functionalized as described previously (Ebner et al., 2007). Briefly, AFM probes were coated with flexible acetal-PEG-NHS spacers (Gruber Lab, Institute of Biophysics, Linz, Austria). In the next step, recombinant Dsg3-Fc produced by Chinese hamster ovary cells as described earlier (Spindler et al., 2009) were coupled to the free acetal bearing end of the PEG spacer. Living keratinocytes were investigated in 1.8 mM $Ca^{2+}$ DMEM medium at 37°C. In order to record the correct unbinding forces during the experiments, the spring constants of the cantilevers was determined using the thermal noise method as described earlier (Hutter and Bechhoefer, 1993). After acquisition of a topographical overview of the scanning area, sampling areas of 3 µm x 3 µm over the cell nucleus and 7.5 µm x 2.5 µm spanning the border of two neighboring cells were selected and force-distance-curves (FDC) were acquired in a pixel wise manner in the force mapping mode. For each FDC the tip of the cantilever was brought into contact with the cell surface at a speed of 10 µm/s until a defined indentation force was reached (0.2 nN, hold for 0.1 s). The tip was then retracted at 10 µm/s for 2 µm and the resulting FDC was recorded. FDCs were analyzed using the JPKSPM Data Processing software (Version 6, JPK Instruments, Berlin, Germany). For each FDC it was determined if a binding event had occurred, identifiable by a jump in the force while the tip was retracting from the surface. The height of the step equals the unbinding force that was necessary to break the bond between a Dsg3 molecule coupled to the tip and a corresponding binding partner on the cell membrane. The position corresponding to the unbinding event (step position) represents the perpendicular distance of the AFM head to the cell surface. Based on the slope of the FDC just before the break of the bond, unbinding events were categorized (Fig. 2A) to be either a “bent” unbinding event (slope<-60 µN/m) or a “tether” unbinding event (slope>-60 µN/m). Specific Dsg3-mediated interactions on HaCaT cells were confirmed under same setting by incubation with anti-Dsg3 AK23 (Biozol) at 75 µg/ml for 30 min,

**Calcium Switch Assays**

To evaluate assembly and disassembly of desmosomes, $Ca^{2+}$ switch assays were performed. α-adducin ctrl and ko HaCaT cells were grown to confluency in DMEM containing 1.8 mM $Ca^{2+}$ ("baseline"). To induce desmosome disassembly, medium was replaced with DMEM containing 5 mM EGTA (VWR) for 60 min ("depletion"). To induce desmosome assembly after depletion, 5 mM EGTA medium was exchanged with DMEM containing 1.8 mM $Ca^{2+}$ for indicated times ("repletion").

**Laminar Flow Assay**

Cells were seeded in 6 channel µ-slides (Ibidi) and grown to confluency. The chamber was placed in an incubator with 37°C and 5% $CO_2$ atmosphere and connected to a tube system where a peristaltic pump (Ismatec, Wertheim, Germany) sucked in culture medium from a reservoir, transported it through the channel with the cells and pushed it finally back into the reservoir where it could get in contact with the $CO_2$ atmosphere. "No flow" channels were left open to the $CO_2$ atmosphere, without attachment to the tube system. The pump was set to produce a force of 6.69 dyn/$cm^2$ in the channels and flow conditions were applied for 24 h.

**Cyclic Stretch Assay**

To stretch cells laterally, they were seeded on 6 well plates with a flexible silicone bottom coated with collagen I (Bioflex, Flexcell® International Corporation, Burlington, NC, USA). After they reached confluency, the plate was attached to the stretch apparatus (Fothy I, CLS cell line service, Eppelheim, Germany) which applied negative and positive pressure to the membrane at 1 Hz. Cells were stretched for 24 h, non-stretched membranes were used as control conditions. After stretch, cells were subjected to either fixation and immunostaining or surface biotinylation assays as described above.

### Transmission electron microscopy

Cells were grown to confluency on coverslips and fixed by adding 37°C 4% PFA + 5% glutaraldehyde (GA) in 0.1 M PIPES with 2 mM $CaCl_2$ in equal volume to the medium. After 15 min, fixative was replaced by 2% PFA + 2.5% GA in 0.1 M PIPES with 2 mM $CaCl_2$ for 2 h at room temperature followed by 16 h at 4°C. They were subsequently washed three times with cold 0.1 M PIPES with 2 mM $CaCl_2$. Fixed cells were rinsed first in PIPES buffer (Sigma Aldrich) and then once in sodium cacodylate buffer (0.1 M, pH7.3) for 10 min. After two additional washes in 0.1 M sodium cacodylate buffer, cells were post-fixed in 1% osmium tetroxide (Electron Microscopy Sciences, Hatfield, PA, USA), 0.8% potassium ferracyanide (Electron Microscopy Sciences) in 0.1 M sodium cacodylate buffer for 1 h at 4°C. Coverslips were rinsed several times in 0.1 M sodium cacodylate buffer and ultrapure distilled water, en block stained with 1% aqueous uranyl acetate (Electron Microscopy Sciences) for 1 h at 4°C in the dark. Cells were then dehydrated in an ethanol series (30%, 50%, 75%, 95% and 100%) at 4°C. After three changes of absolute ethanol, samples were washed in acetone and finally embedded in a mixture of resin/acetone first and then in pure EPON 812 resin (Electron Microscopy Sciences). Coverslips were placed cell-side down on BEEM capsules (Electron Microscopy Sciences) filled with EPON. Embedding was carried out in an oven at 60°C for 48 h. After complete polymerization, coverslips were removed from EPON block using the nitrogen-hot water method. During the removal of the coverslip from the EPON block cells were transferred from the coverslip to the block surface. 70 nm thin serial sections were cut with a diamond knife and placed on formvar-carbon coated copper slot grids, stained with uranyl acetate and Reynolds's lead citrate, and observed in a FEI Tecnai T12 spirit Transmission Electron Microscope (Thermo Fisher Scientific) operating at 80 kV. Images were recorded using a CCD Veleta digital camera.

### Image Analysis

For image analysis, Image J (NIH) was used as well as ICY software (de Chaumont et al., 2012) both with standard functions and plug-ins. To combine and present images Adobe Photoshop CC 2017 (Adobe, San José, CA, USA) and Adobe Illustrator CC 2017 (Adobe) were used.

### Statistics

Statistical computations were performed using Prism 8 (GraphPad Software, La Jolla, CA, USA). To compare two data sets a 2-tailed unpaired Student's t test was performed, for more than two data sets a one-way ANOVA followed by Bonferroni correction was used. Statistical significance was assumed at $p<0.05$. Data are shown as mean ± standard error of the mean.

### SUPPLEMENTARY FIGURES

**Sup. Fig. S1**

(**A**) Western blot of whole-cell lysates of different α-adducin ctrl and ko HaCaT cell lines probed for various adhesion molecules (left) and densitometric quantification (right). n=4, error bars represent SEM. (**B**) Representative immunostainings of α-adducin ctrl2 and ko2 HaCaT cells stained for α-adducin and F-actin. DAPI is used to stain the nucleus. Scale bar: 10 µm. Color code for merged panel: nucleus blue, F-actin green and α-adducin red. (**C**) Western blot of whole-cell lysates of different MEK cell lines from α-adducin wt and ko mice probed for various adhesion molecules (left) and corresponding quantification (right). n=3, error bars represent SEM. (**D**) Representative HE stainings of murine skin biopsies. Scale bar: 100 µm (left). Quantification of epidermal thickness in HE stainings of murine skin biopsies (right). n=9; each dot represents one animal from 4 litters in total, error bars represent SEM. (**E**) Representative immunostainings (left) and corresponding quantification (right) of murine skin biopsies stained for Dsg3. Scale bar: 50 µm. n=4, each dot represents the mean of ≥4 sections of one mouse, error bars represent SEM. (**F**) Quantification of desmosome length in electron microscopic images of confluent α-adducin ctrl and ko HaCaT cells. n≥15 cells, error bars represent SEM.

**Sup. Fig. S2**

(**A**) Interaction probability of Dsg3-functionalized cantilevers on HaCaT ctrl cells under control conditions and with incubation of the monoclonal anti-Dsg3-antibody AK23. n=5, each dot represents the mean value of >1000 force-distance-curves from one independent experiment, *

$p<0.05$ vs. control, error bars represent SEM. (**B**) Representative immunostainings of α-adducin ctrl and ko HaCaT cells stained for Dsg3 after 1 h of $Ca^{2+}$ depletion (left). Scale bar: 10 µm. Scale bar zoom: 5 µm. Quantification of the irregularity score of control and depletion conditions in immunostainings for Dsg3 in ctrl and ko HaCaT cells. n>3, each dot represents the mean of 9 cells of one independent experiment, * $p<0.05$, error bars represent SEM (right).

**Sup. Fig. S3**

(**A**) Representative trajectories of Dsg3-GFP containing vesicles in α-adducin ctrl and ko HaCaT cells. Trajectories are rotated and aligned in a way that from left to right they represent the movement from the intercellular space towards the cell membrane. The duration of each trajectory is 790 s. Scale bar in x and y direction: 2 µm. (**B**) Quantification of average speed (left) and average linearity (right) of the trajectories of Dsg3-GFP containing vesicles in α-adducin ctrl and ko HaCaT cells. n>3, each dot represents the mean of >100 trajectories of at least one cell of one independent experiment, * $p<0.05$ vs. ctrl, error bars represent SEM. (**C**) Representative Western blot of surface biotinylation assay of α-adducin ctrl HaCaT cells. n=3.

**Sup. Fig. S4**

(**A**) Representative super resolution images of apical cell portions of α-adducin ctrl and ko HaCaT cells of immunostainings stained for Dsg3 (left panels). After background subtraction a mask was generated showing clusters of Dsg3 (right panels). Scale bar: 5 µm. (**B**) Quantification of average Dsg3 cluster size (left) cluster intensity (middle) and cluster density (right) from super resolution images of apical cell portions of α-adducin ctrl and ko HaCaT cells of immunostainings stained for

Dsg3. n=3, each dot represents the mean of >5 cells of one independent experiment, error bars represent SEM.

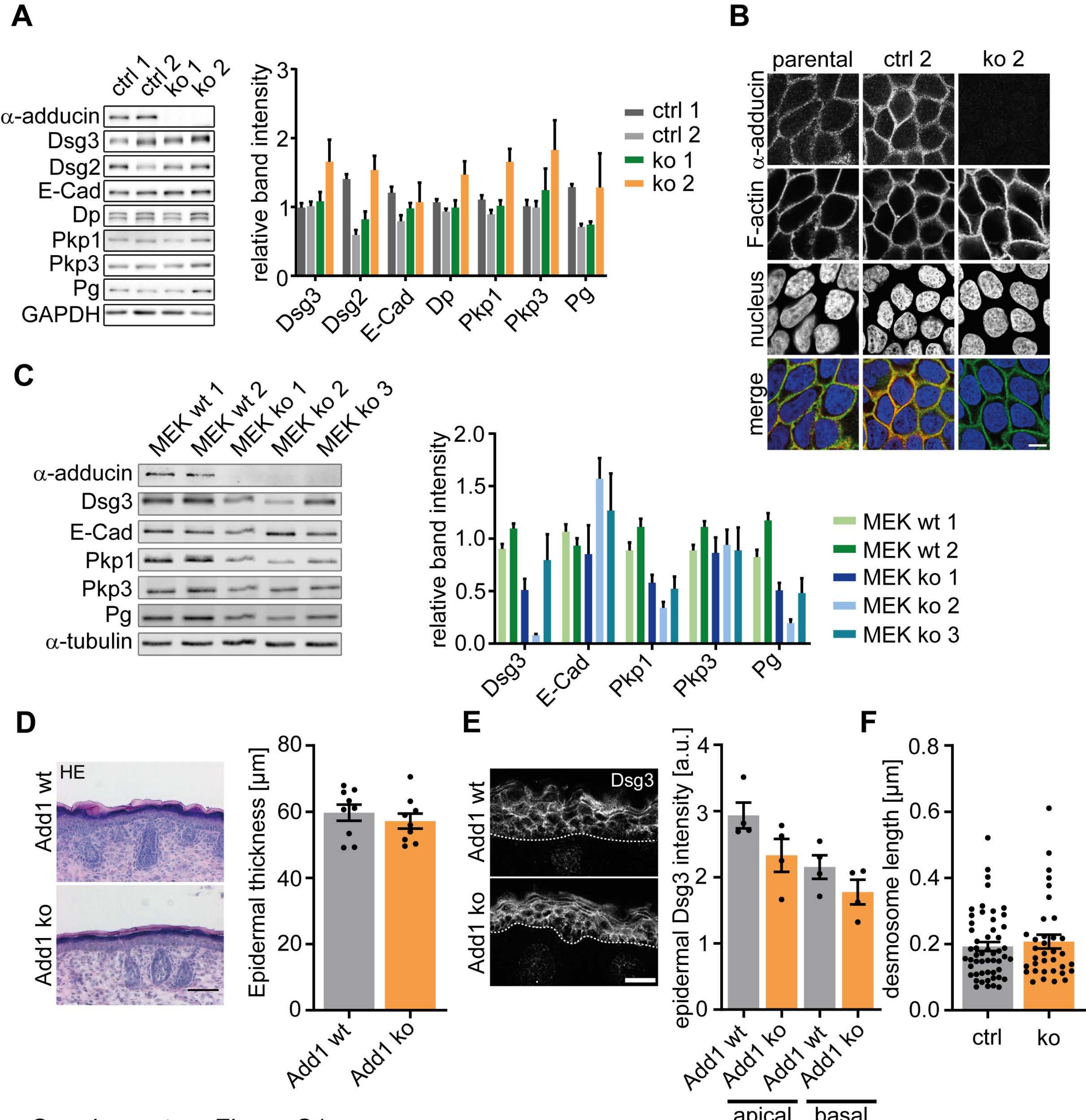


Supplementary Figure S1

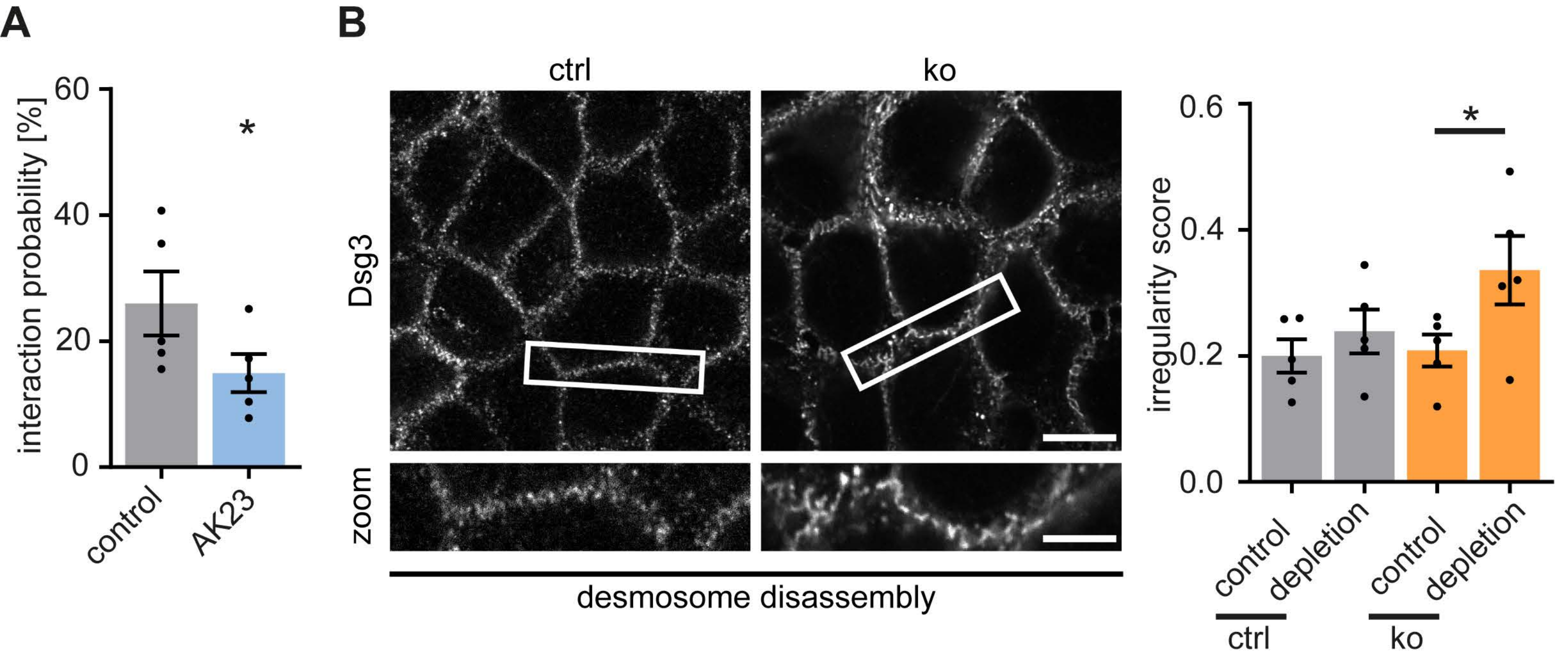


Supplementary Figure S2

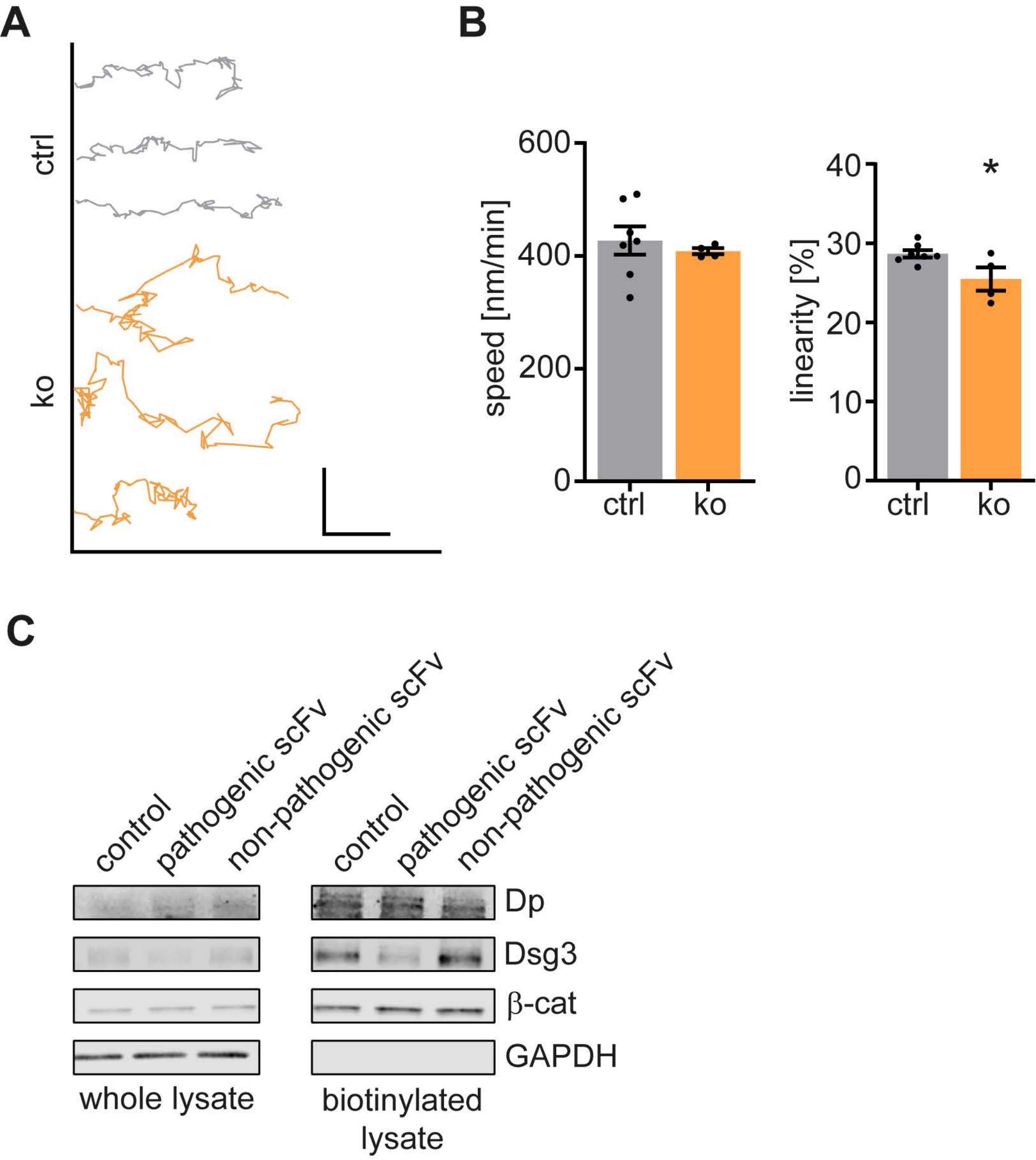


Supplementary Figure S3

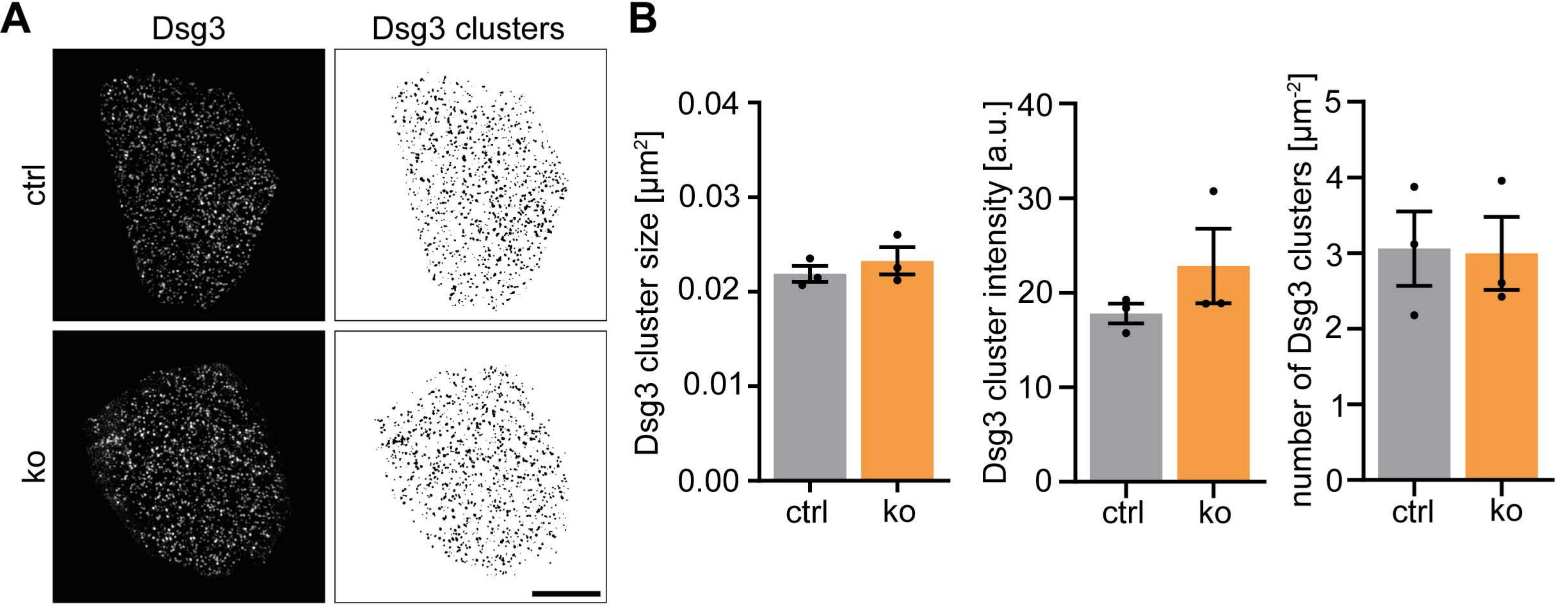


Supplementary Figure S4
